# Adapterama IV: Sequence Capture of Dual-digest RADseq Libraries with Identifiable Duplicates (RADcap)

**DOI:** 10.1101/044651

**Authors:** Sandra L. Hoffberg, Troy J. Kieran, Julian M. Catchen, Alison Devault, Brant C. Faircloth, Rodney Mauricio, Travis C Glenn

## Abstract

Molecular ecologists seek to genotype hundreds to thousands of loci from hundreds to thousands of individuals at minimal cost per sample. Current methods such as restriction site associated DNA sequencing (RADseq) and sequence capture are constrained by costs associated with inefficient use of sequencing data and sample preparation, respectively. Here, we demonstrate RADcap, an approach that combines the major benefits of RADseq (low cost with specific start positions) with those of sequence capture (repeatable sequencing of specific loci) to significantly increase efficiency and reduce costs relative to current approaches. The RADcap approach uses a new version of dual-digest RADseq (3RAD) to identify candidate SNP loci for capture bait design, and subsequently uses custom sequence capture baits to consistently enrich candidate SNP loci across many individuals. We combined this approach with a new library preparation method for identifying and removing PCR duplicates from 3RAD libraries, which allows researchers to process RADseq data using traditional pipelines, and we tested the RADcap method by genotyping sets of 96 to 384 *Wisteria* plants. Our results demonstrate that our RADcap method: 1) can methodologically reduce (to <5%) and computationally remove PCR duplicate reads from data; (2) achieves 80-90% reads-on-target in 11 of 12 enrichments; (3) returns consistent coverage (≥4x) across >90% of individuals at up to 99.9% of the targeted loci; (4) produces consistently high occupancy matrices of genotypes across hundreds of individuals; and (5) is inexpensive, with reagent and sequencing costs totaling <$6/sample and adapter and primer costs of only a few hundred dollars.

## Introduction

Massively parallel sequencing is changing molecular ecology and other life science disciplines (Rogers & Venter 2005; Tautz *et al*. 2010). While the costs of whole genome sequencing and genome resequencing have declined, the time investment, cost, and computational complexity of genome assembly and genome resequencing remain significant drawbacks. Fortunately, many biological hypotheses can be tested with smaller samples of the genome that collect data from several hundred to several thousand variable loci (Cariou *et al*. 2013; Pante *et al*. 2015) rather than requiring the millions of variable sites identified during genome sequencing/re-sequencing workflows. Although genome reduction techniques that collect data from hundreds or thousands of loci are an appealing and inexpensive proxy for whole genome resequencing, the matter of how best to collect genotypes from hundreds or thousands of loci across hundreds or thousands of individuals remains. Thus, researchers still face the decades-old dilemma of choosing among methods that make trade-offs in the number and kind of data (loci) collected versus the number of individuals surveyed.

Genome reduction techniques fall into a broad class of so-called “reduced representation” approaches, and these methods are meant to collect data from a small and repeatable fraction of the genome across a population of individuals - enabling the population under study to be compared at identical loci without sequencing the entire genome (Altshuler *et al*. 2000; Novaes *et al*. 2008; Wiedmann *et al*. 2008). Two general types of reduced representation library approaches for massively parallel sequencing are widely used – sequence capture (Gnirke *et al*. 2009; Okou *et al*. 2007) and restriction-site associated DNA sequencing (RADseq; Baird *et al*. 2008; Davey & Blaxter 2010; Davey *et al*. 2011; Miller *et al*. 2007; Peterson *et al*. 2012).

Although both methods have advantages and disadvantages (Harvey *et al*. 2013), neither is entirely capable of achieving a primary goal of many population genetic studies: consistently obtaining a set of hundreds or thousands of putatively unlinked single nucleotide polymorphisms (SNPs) from hundreds to thousands of individuals at low cost (e.g., <$10/sample).

Sequence capture is a powerful technique that combines a custom set of long, biotinylated, oligonucleotide baits with in-solution hybridization to target and enrich any number of genomic regions of varying size (Gnirke *et al*. 2009), from entire chromosomal segments (Cao *et al*. 2013; Pröll *et al*. 2011; Rabenstein *et al*. 2015) to sets of smaller loci containing SNPs (Kenny *et al*. 2010; Saintenac *et al*. 2011). Sequence capture requires some form of prior sequence information to design capture baits (Gnirke *et al*. 2009), which can be a challenge for certain non-model species and marker types. Several groups have designed bait sets that target conserved sequences including ultraconserved elements (UCEs; Faircloth *et al*. 2012), anchor regions (Lemmon *et al*. 2012), and exons (Bi *et al*. 2012), which allow sets of baits to be used across many species (e.g., at the level of taxonomic class; Li *et al*. 2013). Although useful for many situations, sequence capture is constrained by high library preparation costs, expensive baits, randomness of where the collected sequences start and stop, and off-target sequence reads (Harvey *et al*. 2013). Although off-target reads can be beneficial (Raposo do Amaral *et al*. 2015), researchers must account for such reads to achieve the desired sequencing coverage (e.g., if 50% of reads are off-target, researchers must obtain twice as many reads to yield the same coverage as when all reads are on-target). Conversely, the randomness inherent in the beginning and ending positions of the collected sequences enables the removal of PCR duplicates and probabilistic variant calling methods, which are widely used in genome resequencing research.

RADseq methods reduce the genome by sequencing thousands to hundreds of thousands of DNA fragments that are located near restriction enzyme cut sites (Baird *et al*. 2008; Davey *et al*. 2011; Miller *et al*. 2007). Many RADseq derivatives have been developed (Andrews *et al*. 2016), and throughout this manuscript, we will use the term “traditional RADseq” to mean methods where one end of the sequenced DNA insert derives from a restriction-site and the other end is randomly sheared (Baird *et al*. 2008; Davey *et al*. 2011; Miller *et al*. 2007), whereas we will use “RADseq” to generically refer to any of the derivative forms of RADseq. Dual-digest RADseq (ddRAD; Peterson *et al*. 2012) methods, including our 3RAD variant (Glenn *et al*. 2016b; Graham *et al*. 2015), sequence DNA inserts that fall precisely between two restriction enzyme cut sites, giving both ends of the sequenced DNA precise start and stop positions.

Sequencing these ddRAD-type libraries is particularly efficient when compared to libraries derived from sequence capture or traditional RADseq approaches because sequencing reads pile-up on the ends of the loci, boosting coverage and efficiently increasing the accuracy of downstream SNP calling (Fountain *et al*. 2016). Compared to sequence capture, RADseq methods generally have lower library preparation costs and do not explicitly require existing genomic information from the taxa of interest.

The primary disadvantages of RADseq (see Mastretta-Yanes *et al*. (2015) for review) include sequencing many monomorphic loci and stochastic variation (mutation and methylation) at the restriction enzyme cut sites that produce sparse genotype matrices. Overall quality and utility of RADseq data sets can also be affected by abiotic factors such as the molecular and bioinformatic protocols used to generate the RADseq data. For example, ddRAD loci may be lost due to imprecise size selection methods; errors within loci may be introduced by low-quality reagents; and/or PCR bias may preferentially amplify smaller fragments, GC rich regions (Puritz *et al*. 2014), or one allele over another (Casbon *et al*. 2011).

Particularly problematic are errors introduced to RADseq libraries during PCR. Incorporation errors early during the PCR reaction can be amplified to high coverage as PCR proceeds (Tin *et al*. 2015), and PCR duplication of loci can give false confidence in the accuracy of downstream variant calls. For example, many RADseq processing pipelines use coverage to validate the accuracy of SNP calls and PCR duplicates can comprise 20-90% of reads in RADseq libraries (Ali *et al*. 2015; Andrews *et al*. 2014; Schweyen *et al*. 2014; Tin *et al*. 2015); therefore, duplicate reads can produce inflated coverage across loci, resulting in falsely high confidence assigned to the genotypes obtained (Casbon *et al*. 2011; Schweyen *et al*. 2014; Tin *et al*. 2015).

The traditional approach for distinguishing duplicates in standard genomic libraries, which are randomly sheared on both ends, and traditional RADseq libraries, which are randomly sheared on one end, is to identify duplicate reads as those having identical start and stop positions when aligned to a reference sequence. However, this technique cannot be applied to ddRAD-type approaches, where all sequence reads from each RAD locus are identical (Andrews *et al*. 2014), and it is thought that using less template DNA can exacerbate the problem of read duplication (Casbon *et al*. 2011). Single molecule tagging has been employed to identify and remove PCR duplicates in a variety of approaches (Hiatt *et al*. 2013; Jabara *et al*. 2011; Kivioja *et al*. 2012; Miner *et al*. 2004; Shiroguchi *et al*. 2012; Smith *et al*. 2014), including deep sequencing of limited input DNA. Recently, this approach has been employed in RADseq and ddRAD experiments by incorporating degenerate bases in adapters (Casbon *et al*. 2011; Schweyen *et al*. 2014; Tin *et al*. 2015), but all of the methods developed, to date, have some limitations in their general implementation (see Discussion for details). Single molecule tagging approaches that are easy to implement and have high power to distinguish PCR duplicates are still needed.

Here, we introduce RADcap, a novel method that combines the benefits of single-molecule tagging with 3RAD and sequence capture to collect a consistent and repeatable sample of hundreds of loci across hundreds of individuals, remove PCR duplicates from the resulting data, and call SNPs using a probabilistic base-calling pipeline (GATK; DePristo *et al*. 2011; McKenna *et al*. 2010). The RADcap workflow begins with a pilot experiment using 3RAD to collect genetic information from a small sample of individuals. After processing the resulting sequence reads using Stacks (Catchen *et al*. 2013; Catchen *et al*. 2011) to identify variable RAD loci, the workflow proceeds by designing a set of biotinylated ssRNA baits targeting a subset of the variable RAD loci, and enriching the targeted loci from a pool of DNA libraries prepared using our inexpensive 3RAD library preparation process. To ameliorate the problem of false confidence in genotype calls bolstered by PCR duplicates, the RADcap approach incorporates a random 8nt sequence tag in place of the iTru5 primer index (Figures 1 and 2) to each library molecule, which allows researchers to distinguish PCR duplicates from unique template molecules during post-processing of the sequence data. Finally, following a GATK workflow, we created a RADcap data processing package, which calls SNPs in the duplicate-free reads using a “radnome” (those RAD loci we targeted with capture baits) as a reference sequence. We empirically tested the RADcap method by measuring genetic diversity of 96 samples of *Wisteria* across an urban center.

**Figure 1:**
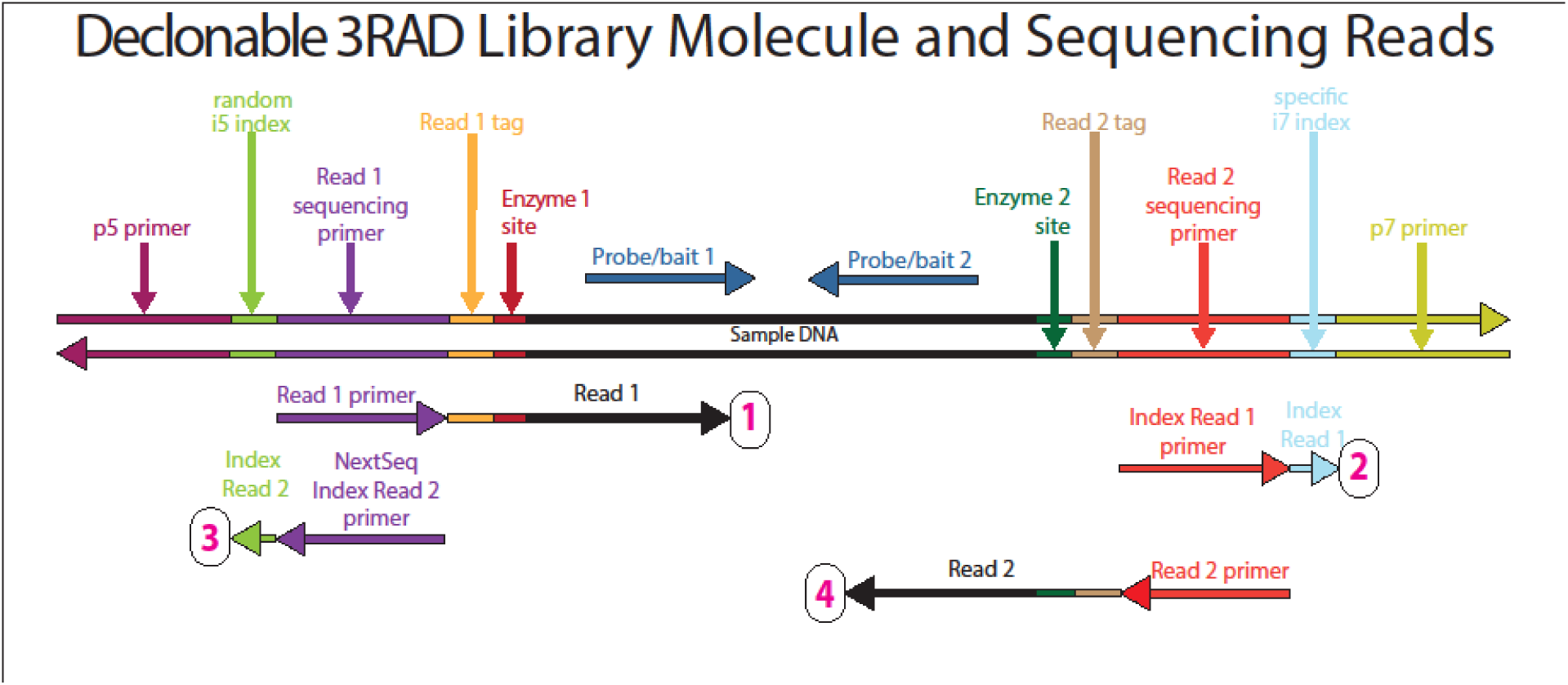
Sequencing reads that can be obtained from full length 3RAD library molecules with iTru5-8N sequence tags. The top double stranded molecule shows a 3RAD library molecule prepared as described in the text. The color-scheme follows those of Glenn *et al*. (2016a; 2016b; 2016c) and Figure 2. The horizontal arrows above the text indicate positions on baits. The horizontal arrows beneath the library molecule indicate Illumina sequencing primers (binding to the complementary strand of the library molecules). The tip of the arrowhead indicates the 3’ end of the primer and the direction of elongation for sequencing. Four sequencing reads are shown for each library prepared molecule, with one read for each index and each strand of the genomic DNA, including internal indexes. Reads are arranged 1 to 4 (numbered in magenta) from top to bottom, respectively. The arrow immediately 3’ of the primers, indicates the data that are obtained from that primer.

**Figure 2:**
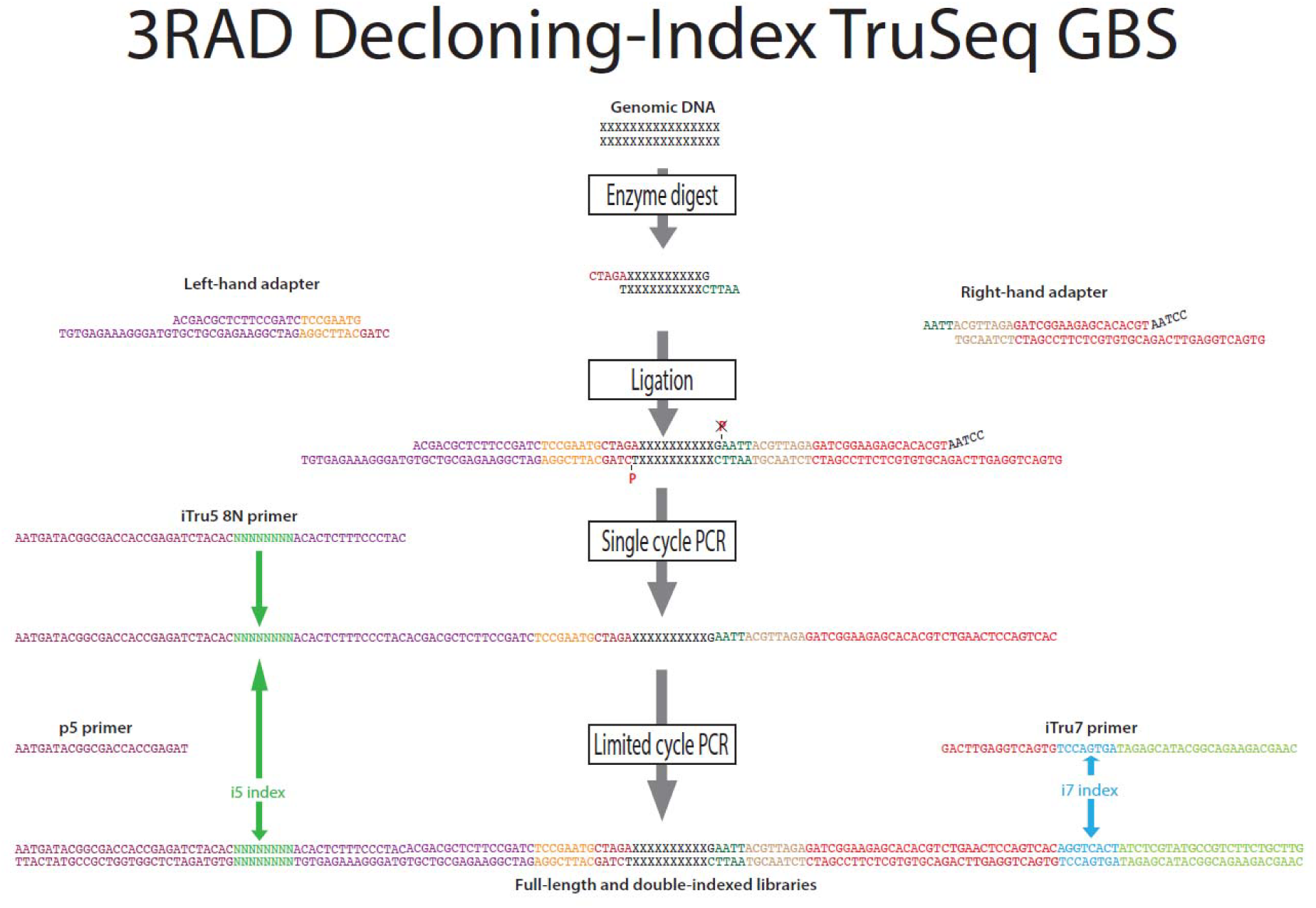
The components of the library molecule added in different steps of the protocol and the sequence of the ends of the molecules. Genomic DNA is digested with enzymes that leave enzyme-specific sticky ends, to which we ligate adapters. The left hand adapter is comprised of four bases that bind to the XbaI restriction site overhang (dark red), a sample-specific internal sequence tag, used to identify the sample (orange) and a Read 1 sequencing primer that is partially single stranded to facilitate annealing of the iTru5 primer (purple). The right-hand adapter is a y-yoke adapter composed of the four bases that bind to the EcoRI restriction site overhang (dark green), a sample-specific internal sequence tag (tan), and the Read 2 sequencing primer (red). During the single cycle PCR, the iTru5 primer is added to the library: the partial library is denatured, the primer anneals to the Read 1 sequencing primer overhang (purple), and extends, thereby adding the degenerate barcode with 8 N bases (green), and the P5 primer (maroon) which anneals to the Illumina flowcell. After cleaning up the reaction, a limited cycle PCR is performed to add the iTru7 primer, comprised of the Read 2 sequencing primer (red) which anneals to the single stranded adapter added earlier, a sample-specific barcode (blue), and P7 primer (light green) which anneals to the Illumina flowcell.

## Methods

### Study Species and Experimental Design

Previous research on *Wisteria* in the southeastern United States (Trusty et al. 2007; Trusty et al. 2008) and within Athens, GA (Glenn *et al*. 2016c) has shown that most *Wisteria* plants are hybrids of *W. sinensis* and *W. floribunda*. Both species were introduced to the United States in the early 1800s (Wilson 1916; Wyman 1949) as ornamental plants and both species reproduce sexually and vegetatively (Valder 1995). While currently available genetic markers can distinguish species, there are no markers available with enough resolution to distinguish among individuals from the same population. Understanding the population genetics of this invasive plant requires many more markers, and is crucial to understanding how it is spreading. We use SNPs to estimate the genetic diversity of 96 samples across Athens, Georgia, USA. We compare these estimates to estimates of genetic diversity obtained from SNPs from the same loci in the same samples, that were prepared via the more traditional 3RAD instead of RADcap.

### 3RAD SNP Discovery for Bait Design

Because sequence capture uses baits designed from pre-existing sequence information, we collected these data using a pilot 3RAD study of four individual *Wisteria* plants: three samples collected around Athens, GA (wist69-3, wist124-1, and wist276-4) and one sample collected from greenhouse-grown seedlings (Wmat9-7-P5-S1). We prepared samples using 3RAD (Glenn *et al*. 2016b; Graham *et al*. 2015), which we summarize below and explain, in detail, in the Supplemental Methods. We added short forward and reverse adapters containing a unique tag combination to extracted DNA from each of the four samples, and we performed a restriction digest of this solution using XbaI, EcoRI-HF, and NheI-HF. Following initial digestion, we added T4 DNA ligase to the digested DNA without disabling the restriction enzymes, and we cycled temperatures to sequentially promote ligation of adapters followed by digestion of chimeras and dimers. We cleaned the resulting reactions with NaCl-PEG diluted Speedbeads (Rohland & Reich 2012), and we completed the adapter sequences using PCR with iTru5 and iTru7 primers (see Glenn *et al*. (2016a) for details about primers). We pooled the resulting libraries, size-selected fragments of 550bp (+/- 10%) with a PippenPrep (Sage Science, Inc.), and performed a final round of low-cycle PCR-recovery using the P5 and P7 primers to increase the concentration of fragments in the desired size range. We sequenced samples on an Illumina NextSeq v2 300 cycle kit to obtain paired-end 150nt (PE150) reads (Figure 1).

We used the *process_radtags* program in Stacks v1.29 (Catchen *et al*. 2013; Catchen *et al*. 2011) to clean and demultiplex the resulting sequence data. We “rescued” sequence tags and RAD-tags within 2bp of their expected sequence; otherwise, we removed reads with an uncalled base or containing the wrong adapter or wrong cut site. Because our 3RAD adapter sequences vary in length, and because Stacks requires all reads to be the same length, we truncated reads to 140bp, removing 0-3 bases of sequence per read, in *process_radtags*. We ran the Stacks pipeline with the following modifications: in the *ustacks* program, we removed highly repetitive stacks, we used the deleveraging algorithm, and we set the maximum distance between stacks (M) to 3; in the *cstacks* program, we set the number of mismatches allowed between sample tags when generating the catalog (n) to 4; in the program *populations,* we required at least 3 individuals to have reads to retain a given locus (r), and we set the minimum stack depth required for individuals at a locus (m) to 3. We output the full sequence from each allele identified across our pilot samples in FASTA format. We selected loci that were polymorphic, but had less than five SNPs across both paired end reads, and that were present in three or four of the samples, which resulted in 1740 paired reads (candidate loci) for bait design.

### Bait Design and Synthesis

We selected bait sequences to minimize target redundancy and bait-to-bait hybridization, which can compromise the synthesis of ssRNA baits as well as the target capture hybridization reaction. To perform these steps, we subjected sequences to self-analysis using BLAST 2.2.19 (*filter query sequence = false, word size = 11, e-value = 1e-13, number of sequences to show alignments for = 2,000;* Boratyn *et al*. 2013). We discarded any locus with one or both sequences having a BLAST hit of at least 140bp to another sequence (682 loci). Next, we subjected sequences to a same-strand self-analysis in BLAST (as above, *query strand = bottom*). We discarded 94 additional loci in which one or both paired sequences had a BLAST hit to another sequence, leaving 964 loci. Then, we designed two sets of 90mer baits targeting the remaining 964 loci. In the first set, we chose a single bait from both paired sequences for every locus, and we positioned baits to start at the 20th base of their parent sequences (creating 1,928 *Wisteria* baits; Supplementary File: Wist-Probes-Set1.fasta). In the second set, we added additional baits from both sequences corresponding to a random subset of 200 loci (creating 400 additional baits; Supplementary File: Wist-Probes-Set2-SUBSET-400.fasta), and we positioned these baits to start at the 40th base of their parent sequences. The two sets produced a total of 2,328 baits targeting *Wisteria* library molecules. To reduce synthesis costs, we combined this bait set design with a similar number of baits designed in the same way for another species (*Pueraria montana* var. *lobata,* kudzu). We subjected the bait sequences for both species to a final same-strand self-analysis using BLAST (same process as above), and we did not find evidence of additional bait-to-bait hybridization. Before bait synthesis and because MYbaits^®^ cannot be synthesized with a mixture of bases, we replaced any variable positions in any bait sequence with a random candidate base, and we replaced all unknown (“N”) positions with a thymine. We created a custom set of biotinylated RNA baits by having them synthesized as a MYbaits-1 kit (MYcroarray, Ann Arbor, MI, USA).

### Sample Library Preparation and Description of Treatments

We provide detailed sample collection and sample preparation methods in the Supplemental Methods. Briefly, we randomly arranged 192 of the 203 greenhouse-grown *Wisteria* samples in 2 plates (RADcap_Plate1 and RADcap_Plate2; Supplementary Tables 1 and 2). We placed the remaining 9 greenhouse-grown samples into a third plate, to which we added randomly selected columns of the DNA in plates 1 and 2 (RADcap_Plate3; Supplementary Table 3). This arrangement allowed us to re-process 133 libraries independently prepared from the same samples, and we used these replicates to compute the amount of missing data between replicate samples that was not caused by genetic variation. Five samples were included twice across plates 1-3, and 41 samples were included three times across plates 1-3. We arranged the 192 samples from wild-collected individuals by DNA concentration in a fourth and fifth plate (RADcap_Plate4 and RADcap_Plate5; Supplementary Table 4 and 5). We normalized DNA concentration, then we digested the plated DNA with XbaI, NheI-HF, and EcoRI-HF in a reaction that included forward and reverse adapters. As before, we added T4 DNA ligase to the digested DNA without disabling the restriction enzymes, and we cycled temperatures to sequentially promote ligation of adapters followed by digestion of chimeras and dimers (Figure 2; see Supplemental Methods for full details). Following adapter ligation, we combined approximately 66% of the ligation volume from each sample in each plate into plate-specific pools, we cleaned each pool with speedbeads, and we re-suspended cleaned pools in 33ul of TLE. For plates 1-4, we split each pool into three aliquots of 20μl, 10μl, and 3μl, and we used these aliquots to test the effect of different PCR conditions on the efficiency of RADcap (described below; Table 1).

#### Single molecule tagging

To tag and track duplicate reads that resulted from the PCR amplification process, we designed a new iTru5 i5 primer (Glenn *et al*. 2016a) that incorporated a random 8 nucleotide sequence tag (i.e., the i5 index sequence was specified as NNNNNNNN when ordering the iTru5-8N primer). This resulted in the synthesis of a mixture of 65,536 iTru5 i5 primers with unique, 8 nucleotide index sequences. In the experimental treatments, below, we incorporated these uniquely tagged iTru5-8N i5 primers to our DNA library constructs using different PCR conditions to determine what methods produce the fewest PCR duplicates.

#### One-primer amplifications

Following adapter ligation and cleaning, we split each 20μl aliquot into two tubes to increase the total PCR volume possible, and we performed a single-cycle, one-primer PCR. Each reaction contained 10μl template DNA and the iTru5-8N primer. Because we amplified each reaction using only one cycle, the primers did not denature from the library molecules and re-anneal to different library molecules. We pooled the two resulting reactions and cleaned them with speedbeads, and we split them into 2 tubes for a 6-cycle PCR where we included the P5 primer and the plate-specific iTru7 primer (Supplementary Table 13). This second reaction completed the library construct, added the plate-specific i7 index sequence to each library construct, and increased the total amount of library available for capture. We called the plates in this treatment RADcap_1cycle_Plate1-4.

#### Two-primer amplifications

For the second aliquots of 10μl, we performed four PCRs for each pooled plate with 2μl template DNA in each. We included both the iTru5-8N primer and the iTru7 primer in each PCR, and we ran PCR for 5 cycles. Because we included the iTru5-8N primer in the PCR reaction for multiple cycles, newly synthesized molecules could receive new i5 tags (Casbon *et al*. 2011) and thus a single template DNA molecule could generate multiple library constructs with unique i5 sequence tags (i.e., this method produced ≤10 undetectable PCR duplicates per template molecule). Because we used libraries in these treatments that were identical to those used above (i.e., one-primer amplification with a single-cycle PCR), this experiment allowed us to determine the effect of low-efficiency first-strand replication and test how additional PCR cycles affect the identification of PCR duplicates and subsequent variant calling. We called the plates in this treatment RADcap_5cycle_Plate1-4.

#### Low-template, one-primer amplifications

We used the 3μl aliquot from plate 1 to determine the effect of low DNA concentrations on PCR duplication and subsequent variant calling. We added the iTru5-8N primer to 3μl of template from plate 1, and we a performed a single-cycle PCR. As we described for the one-primer amplifications above, we cleaned the resulting PCR product and performed another 6-cycle PCR to add the iTru7 primer. We called this treatment RADcap_Low_Template_Plate1.

#### RAD locus capture

Following all final PCRs described above, we pooled the replicate PCRs, cleaned each plate pool with speedbeads, and performed a separate capture hybridization reaction on each pool from each treatment (9 total) according to the MYcroarray MYbaits v3.0 protocol, with a hybridization temperature of 65°C for 21 hours. Following capture, we split each capture into three tubes and amplified loci with the P5 and P7 primers in an 18 cycle PCR. PCRs from each capture were pooled and cleaned with speedbeads. Following PCR recovery and cleanup, we quantified the 9 experimental treatments, and we pooled these libraries with unrelated libraries from other experiments at a ratio that would return 20% of the reads from an Illumina sequencing run. We sequenced the pooled libraries using an Illumina NextSeq High Output v2 150 cycle kit to achieve PE75 reads.

#### Blocking over-enrichment and testing RADcap versus size selection

After sequencing, the data from plates 1-4 included 8 loci with an average coverage 20x higher than other loci in the one-primer treatment, 10x higher than other loci in the two-primer treatment, and 28x higher than other loci in the low-template treatment. To block the over-enrichment of these loci, we designed and ordered 29 custom oligonucleotides (Supplementary Table 6) between 26 and 60 bp long that were complementary to the baits targeting these 8 loci and which had a DNA to RNA T_m_ greater than 70°C. We also optimized the PCR for plate 5, based on the one-primer treatment above, by increasing reaction volumes 3-fold for the PCR reaction to add the iTru5-8N primer and 1.3-fold for the PCR to add the iTru7 primer, and we included the locus-specific bait blockers during the hybridization reaction. To compare the results of capturing RAD loci to those of size selection normally done in 3RAD (and other RADseq protocols), we split plate 5 in half following the one-primer PCR, and we captured loci from one-half of the plate, as described above. We called this treatment RADcap_optimized_Plate5. We size-selected the remaining half of plate 5 samples as described above, for SNP discovery. We called this treatment 3RAD_SizeSelect_Plate5. We pooled these two libraries with unrelated libraries to obtain 7% of the reads on a second Illumina NextSeq run using conditions described above for the other RADcap libraries.

## Data Analysis

### Modification of Stacks software

Stacks (Catchen *et al*. 2013; Catchen *et al*. 2011) had previously been modified to identify the variable-length internal tags that distinguish individual samples in 3RAD data. However, no software program existed to properly identify and remove the PCR duplicates from RADseq data. We developed and implemented new code as part of the *clone_filter* module within Stacks v1.35 to remove PCR duplicates. *Clone_filter* can be used before or after *process_radtags* and can use any combination of inline or index sequence tags, in addition to using read sequences, to reduce duplicated reads to a single representative in the output. Importantly, *clone_filter* does not modify FASTQ headers, allowing repeated use of *process_radtags* and *clone_filter* for read demultiplexing and duplicate removal.

#### Sample demultiplexing, alignment, and SNP calling

After sequencing, we converted BCL files to FASTQ format using bcl2fastq2 v2.16.0.10 (Illumina, Inc.), and we modified the default parameters to create a separate FASTQ file for index reads (Figure 1). We demultiplexed and removed PCR duplicates from the FASTQ data using Stacks v 1.35. First, we demultiplexed reads by i7 tag (Supplementary Table 13) using *process_radtags*. We discarded reads with an uncalled base, reads having low quality (using default settings), or reads having a sequence tag or RAD tag more than 2 bases distant from the expected sequence, and we rescued reads having sequence tags or RAD tags and within 2 bases of the expected sequence. This initial demultiplexing produced paired end files corresponding to each plate in each treatment. We ran *process_radtags* again on each plate of samples, with the same parameters, to demultiplex reads by inner adapter, which produced paired end files for each individual in each plate. Finally, we used the *clone_filter* program to remove any read having the same combination of random i5 tag and insert sequence, which likely represent duplicates created during PCR amplification.

We created a FASTA-formatted “radnome” file that contained the 964 paired sequences from which we designed baits, and we used this file as a reference sequence for read alignment and SNP calling (Supplementary File: wisteria_reference.fasta). Within this FASTA file, paired reads were separate entries given arbitrary locus names, and we inserted 20 Ns between the sequences for Read 1 and Read 2. We aligned RADcap reads to the reference using BWA v 0.7.7 (Li & Durbin 2009) with the mem algorithm and shorter split hits marked as secondary (M), and we called SNPs using an automated pipeline (https://github.com/faircloth-lab/radcap) that incorporates BWA, PICARD, and the open-source GATK-lite package (DePristo *et al*.2011; McKenna *et al*. 2010). Following automated BWA alignment, the pipeline merged individual alignments, re-aligned BAM files around indels, called SNPs and indels, and filtered problematic or low-quality SNP calls from the total set of raw SNP calls to create a passing file of SNPs.

Variant calling is an inherently population-based process in that errors can be distinguished from variants at a specific position by considering that position in all individuals in the population (Bansal *et al*. 2010; Catchen *et al*. 2011; Craig *et al*. 2008). Therefore, the detection and statistical properties of variant genotypes are dependent on how the population under study is sampled, with fewer variant sites recovered with lower statistical support from smaller populations. To mimic this effect of population sampling and to facilitate comparisons among our experimental treatments, we called SNPs in two ways. First, we treated all 384 individuals from plates 1-4 as a single population, and we called SNPs separately in each of the one-primer (n=384 individuals) and two-primer experimental treatments (n=384 individuals). Second, we treated the 96 samples in plate 1 as a single population, and we called SNPs for the plate 1 population in the one-primer, two-primer, and low-template treatments, as well as the plate 5 optimized and size selected treatments (Table 1). After SNP calling, we filtered the resulting VCF files using vcftools v0.1.12b (Danecek *et al*. 2011) to exclude sites with more than 50%, 20%, or 10% missing data (i.e., 50%, 80%, or 90% complete data), and we computed summary statistics across captured loci and variant sites using a program (radcap_summarize_snp_calls.py) from the *radcap* software package.

**Table 1:** Overview of treatments, DNA plates used with each treatment, and how plates were grouped for analyses. The iTru5-8N reaction volume for the size selected and optimized treatment represents the same reactions, as these treatments were split after the single-primer PCR and clean up.

| Treatment | Plate ID's | Cycles with iTru5-8N | iTru5-8N reaction volume (µl) | All PCR duplicates tagged? | Captured? | Analysis groups (plate ID's) |
| --- | --- | --- | --- | --- | --- | --- |
| RADcap_1cycle | 1 - 4 | 1 | 100 | Yes | Yes | 1; 1-4 |
| RADcap_5cycle | 1 - 4 | 5 | 100 | No | Yes | 1; 1-4 |
| RADcap_Low_Template | 1 | 1 | 25 | Yes | Yes | 1 |
| RADcap_optimized | 5 | 1 | 300 | Yes | Yes | 5 |
| 3RAD_SizeSelect | 5 | 1 | 300 | Yes | No | 5 |

#### Efficiency of Random Tagging at the i5 Index Position

Because PCR can be biased by the composition of certain primers, we wanted to estimate how well our iTru5-8N primers were incorporated into our library constructs. Using the FASTQ file of index reads as input, we determined the count of each iTru5-8N sequence tag using FastX v0.0.14 (Gordon & Hannon 2010; Supplementary file: i5_and_coverage_code.md). We plotted the cumulative count of iTru5-8N sequence tags incorporated to DNA libraries for all possible sequence tag combinations, except for iTru5-8N tags that do not return a signal on the NextSeq (GGGGGGGG), those DNA inserts that have no apparent i5 sequence tag (AGATCTCG), and those iTru5 sequence tags in the adapters ligated to other libraries on the sequencing run.

#### RADcap Efficiency and Coverage

We expected sequence capture to be more efficient than size selection and that the resulting data from captured RAD loci would include fewer off-target reads, have higher coverage at target loci, and consistently recover a larger number of target loci from reads. To investigate these parameters, we computed the coverage of each position in each sample from BAM files using SAMtools v1.2 (Li *et al*. 2009; Supplementary file: i5_and_coverage_code.md). For this analysis, we used the BAM files produced directly from BWA to avoid effects of the BAM re-alignment on our coverage computations and because we wanted to assess which loci were present in the dataset (where coverage of loci in the radnome reference was greater than 0), despite being monomorphic or having errors. We report the average coverage for all loci and samples within each plate, normalized by million reads per sample. In order to determine whether the variation in coverage between loci in a treatment decreased in the optimized treatment, we plotted the log transformed coverage of each locus and tested whether the optimized treatment had less variation in log transformed coverage using a one-sided Siegel-Tukey Test for equality in variability with adjusted medians in DescTools (Signorell 2015) in R. We then calculated the average coverage per locus per million reads per sample for loci with at least 4x coverage in plates RADcap_1cycle_Plate1, RADcap_5cycle_Plate1, RADcap_Low_Template_Plate1, RADcap_optimized_Plate5, and 3RAD_SizeSelect_Plate5. As a measure of consistency and to see if the same loci were recovered in each treatment, we identified the loci with at least 4x coverage in 90% of samples from each treatment and determined the loci in common between treatments using VennDiagram (Chen & Boutros 2011) in R. In addition, we plotted the density kernel of the coverage for Read 1 and Read 2 for each of the five treatments, and compared the distributions of coverage between treatments in a one-sided two-sample Kolmogorov–Smirnov test in R.

To determine how many reads were necessary to recover all of the targeted loci at reasonable coverage, we plotted the number of loci at or above 4x coverage and the number of reads for each sample. To get more resolution at lower read numbers, we took the median coverage for all samples at each locus and divided that to get corresponding coverage between 1,000 and 100,000 reads per sample. We plotted the number of loci at or above 4x, 10x, and 20x coverage as a function of the reads per sample.

#### Error Rate and Genetic Diversity of Wisteria

We calculated the frequency of missing data between replicate samples within the one-primer and two-primer treatments by converting VCF output files to Genepop format in PGDSpider v. 2.0.9.1 (Lischer & Excoffier 2012) and counting the number of SNPs at which one sample had a base called while another did not. Since there were no replicate samples within plate 5, we could not assess the amount of missing data. Instead, we compared estimates of genetic diversity of *Wisteria* in plates RADcap_optimized_Plate5 and 3RAD_SizeSelect_Plate5 from 80% filled matrices in GenAlEx v6.502 (Peakall & Smouse 2006; Smouse & Peakall 2012). For each plate, we report the average number of samples genotyped (out of 96) across all loci, the number of alleles identified, the effective number of alleles, Shannon’s Information Index, the observed and expected heterozygosity, and the fixation index, along with standard error estimates for each parameter.

## Results

### Initial 3RAD SNP Discovery for Bait Design

Following SNP discovery using four Wisteria samples, we obtained 1.4 to 2.5 million PE150 reads per sample, and we retained an average of 83.7% of reads after quality filtering.

We identified 31,686 loci placed in the Stacks catalog, 1350 of which were sequenced in all four samples, and 3483 of which were sequenced in three samples. Of the loci recovered in at least three samples, 2573 loci were polymorphic and contained a total of 6531 variant sites. After filtering these loci, there were 1428 putative variants in the 964 loci we used to design our capture baits.

### Random Tagging at the i5 Index Position Allows Removal of PCR Duplicates

For the RADcap samples in plates 1 to 5, we obtained 3-40 million reads per plate (average 17 million), and we retained >94% of reads after quality filtering (Table 2). We incorporated and sequenced all 65,536 of the expected i5 random sequence tags in both of the Illumina NextSeq runs performed to generate our data, and after removing tags from other libraries on the run, and tags indicating no i5 or no signal, we did not observe major sequence dependent biases that affected PCR efficiency of primers containing different random i5 sequence tags (Supplementary Figure 1). All plates from which we collected data using RADcap had a similar percent of reads retained after quality filtering by *process_radtags* in Stacks (Table 2). We retained an average of 68.9% of reads after decloning (range 20.4 to 95.7%; Table 2, Supplementary Tables 7-11), with the most reads retained from the optimized PCR protocol (which we performed on RADcap_optimized_Plate5 and 3RAD_SizeSelect_Plate5) and RADcap_5cycle_Plate3.

**Table 2:** The total reads, percent retained after quality filtering in *process_radtags,* percent retained after decloning with *clone_filter,* percent mapped with BWA mem algorithm for each plate, and the average coverage per million reads sequenced per sample of all loci.

| Plate | Number of reads | % retained after quality filtering | % retained after decloning | % reads that map to reference | Average Coverage |
| --- | --- | --- | --- | --- | --- |
| RADcap_1cycle_Plate1 | 19,397,440 | 94.9 | 25.1 | 79.6 | 712 |
| RADcap_1cycle_Plate2 | 14,703,752 | 95.0 | 20.4 | 85.7 | - |
| RADcap_1cycle_Plate3 | 15,865,294 | 94.8 | 67.2 | 81.5 | - |
| RADcap_1cycle_Plate4 | 17,907,048 | 95.0 | 63.3 | 83.8 | - |
| RADcap_5cycle_Plate1 | 18,045,032 | 95.0 | 83.3 | 84.8 | 830 |
| RADcap_5cycle_Plate2 | 17,968,264 | 95.0 | 86.9 | 85.5 | - |
| RADcap_5cycle_Plate3 | 2,332,154 | 94.1 | 95.7 | 84.3 | - |
| RADcap_5cycle_Plate4 | 18,455,900 | 95.1 | 86.1 | 84.1 | - |
| RADcap_Low_Template_Plate1 | 3,285,096 | 95.3 | 41.0 | 65.8 | 612 |
| RADcap_optimized_Plate5 | 17,929,096 | 95.5 | 94.2 | 90.1 | 764 |
| 3RAD_SizeSelect_Plate5 | 39,543,602 | 95.9 | 94.8 | 15.1 | 142 |

### Optimizing RADcap Efficiency and Coverage

All but one of the capture treatments yielded ≥80% of reads on target (Table 2), while the optimized treatment (RADcap_optimized_Plate5) yielded the highest proportion of reads on target (90%). More traditional 3RAD with size selection (3RAD_SizeSelect_Plate5) yielded 15% of reads on target. Similarly, the optimized and two-primer treatments had the highest average coverage, at 764 and 830 reads per million reads per sample, respectively (Table 2), but we note that the two-primer coverage is inflated with undetected duplicate sequences from multiple rounds of PCR. The one-primer and low-template treatments had slightly lower average coverages, at 712 and 612 reads per million reads per sample, respectively. The size-selected treatment had the lowest average coverage at 142 reads per million reads per sample.

The coverage per locus per million reads was much higher among the RADcap samples than traditional 3RAD size-selected samples (Figure 3). The increased performance of RADcap is also apparent when the coverage per locus is plotted as a density distribution (Supplementary Figure 2). Among RADcap samples, the variation in coverage per locus per million reads sequenced per sample was lower for RADcap_optimized_Plate5 than 3RAD_SizeSelect_Plate5 and RADcap_1cycle_Plate1 (p < 0.0083 in both cases), but the variation in coverage for the RADcap_optimized_Plate5 did not differ from the RADcap_5cycle_Plate1 or the RADcap_Low_Template_Plate1 (p > 0.37 in both cases; Figure 3).

**Figure 3:**
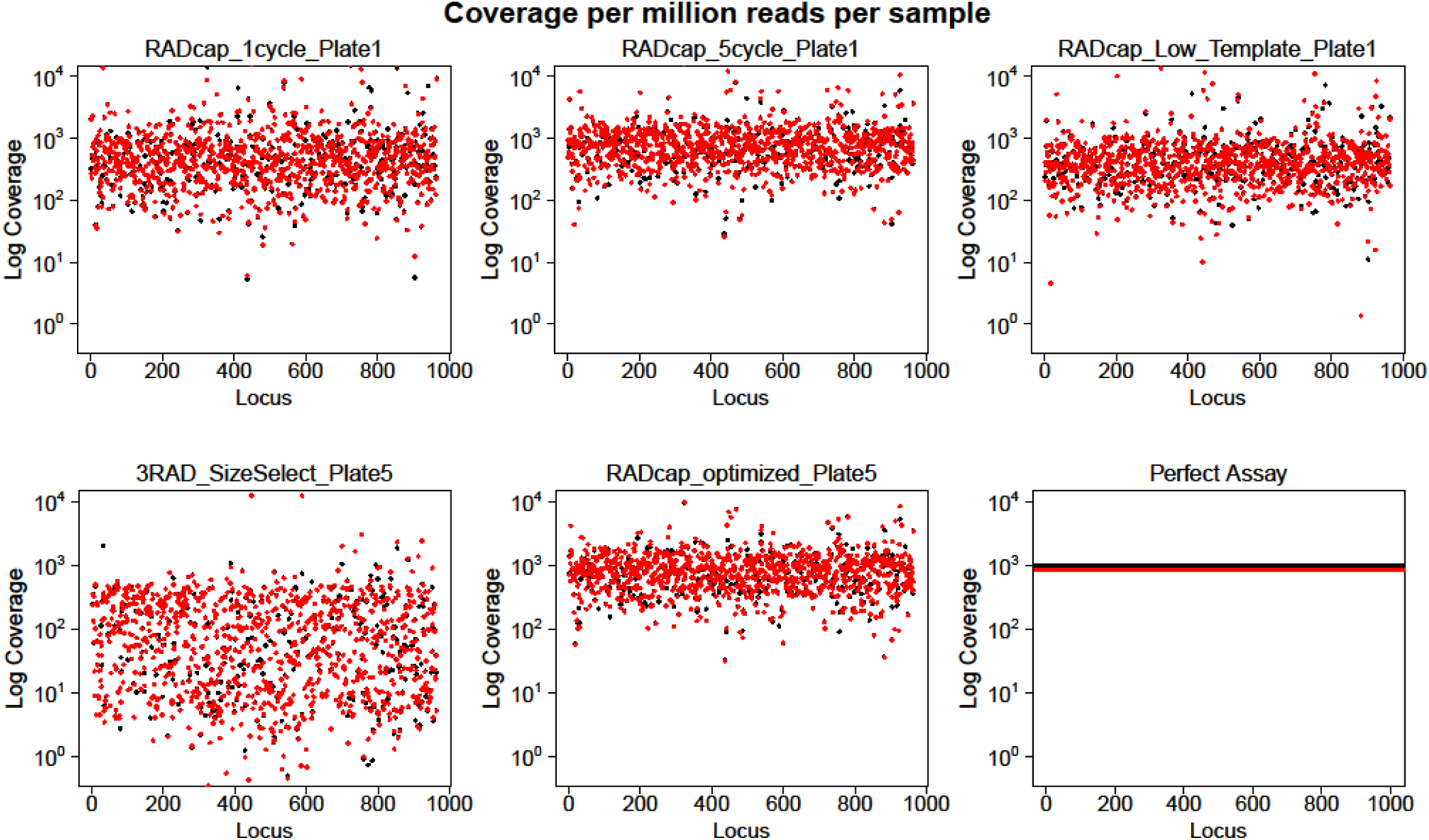
The average coverage (log-transformed) per million reads per sample for 96 samples plotted on a log scale for the y-axis. Coverage for Read 1 is in black and Read 2 is in red. The bottom right panel represents a perfect assay, where all loci have the same coverage of 1000x per million reads per sample. The optimized treatment has a lower variance in coverage across loci than the size-selected or one-primer treatments (p < 0.0083), but variance in coverage was similar to the two-primer or low-template treatments (p > 0.37).

### RADcap Effectively and Consistently Enriched Target Loci and Produces Dense SNP Matrices

We consistently recovered more targeted loci within RADcap treatments than traditional 3RAD with size selection (Figure 5; Supplementary Table 12). Specifically, the optimized treatment performed the best, with 912 loci recovered at 50% matrix occupancy, 880 recovered at 80% occupancy, and 820 recovered at 90% occupancy. The 96 samples analyzed from the one-primer and two-primer treatments performed slightly poorer, with 840 and 874 loci recovered at 50% matrix occupancy, 697 and 823 loci recovered at 80% matrix occupancy, and 552 and 764 loci recovered at 90% matrix occupancy, respectively. Traditional 3RAD with size selection returned 821, 642, and 510 loci at the same levels of matrix occupancy. As expected, RADcap_Low_Template_Plate1 showed the poorest performance, returning 730, 338, and 155 loci at the same levels of occupancy. The number of SNPs called within loci showed similar patterns (Fig 5b), with RADcap_optimized_Plate5 performing better than all other treatments. The effects of population size on the number of SNPs called is apparent in the differences we observed between RADcap_5cycle_Plate1 and RADcap_5cycle_Plate1-4 and RADcap_1cycle_Plate1 and RADcap_1cycle_Plate1-4.

**Figure 5:**
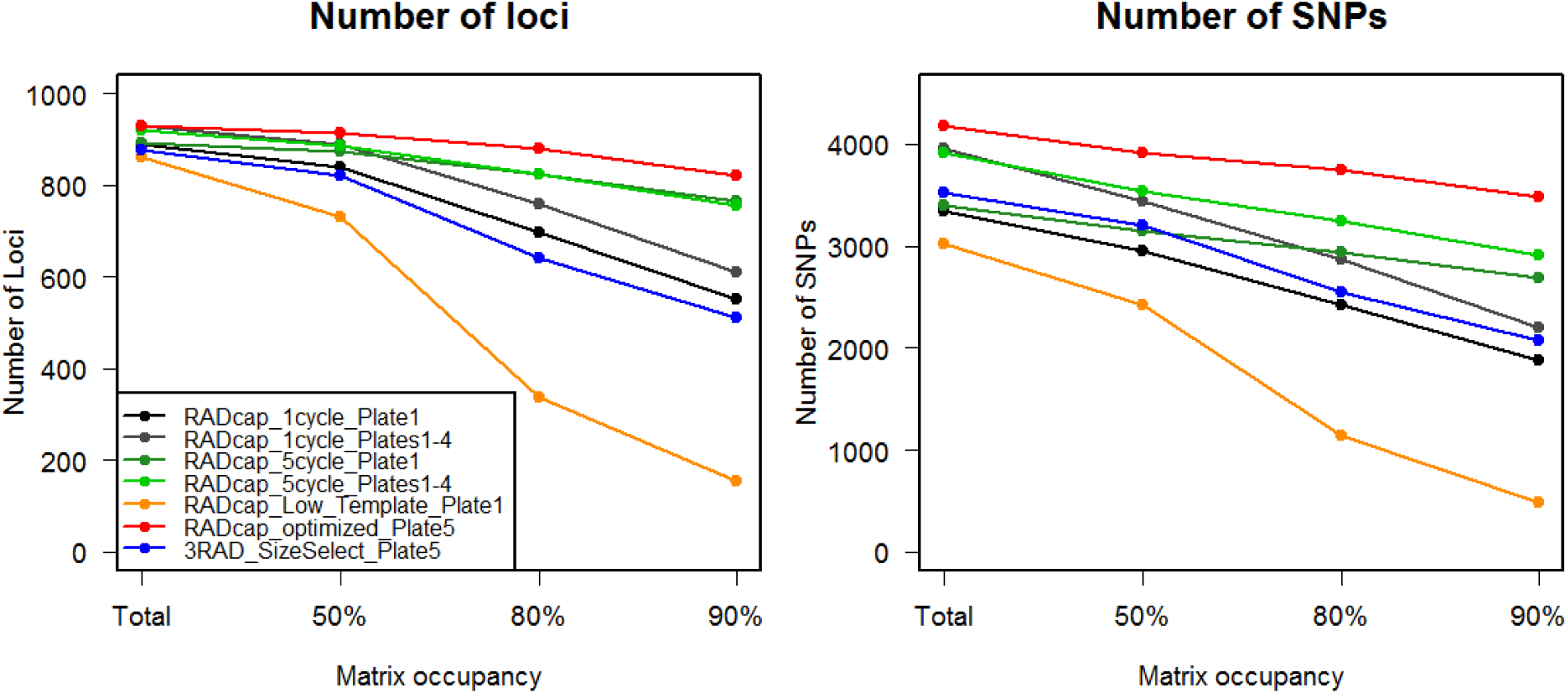
The number of loci and SNPs retained at various levels of matrix occupancy for different treatments, analyzed with GATK. The number of loci and SNPs is the highest and most consistent in the optimized treatment.

Although the effectiveness of enrichment within a plate is one metric of consistency, the more important metric for most researchers is the consistency with which reduced representation approaches collect data across plates or from all individuals in a population. At 90% occupancy for 4x coverage, more than half of the loci (516 of 964; 54%) were shared between all treatments except low-template, an additional 286 loci (30%) were shared among all three RADcap treatments, and 125 loci (13%) were shared among the RADcap_5cycle_Plate1, RADcap_optimized_Plate5, and 3RAD_SizeSelect_Plate5 (Figure 4). Impressively, RADcap_optimized_Plate5 contained data at 4x coverage for 962 of the 964 loci (99.8%; Figure 4). Only 34 loci (3.5%) were present in only two treatments, and only 2 loci (0.02%) were present in a single treatment (Figure 4). Thus, most loci were present in most samples no matter from which treatment they originated.

**Figure 4:**
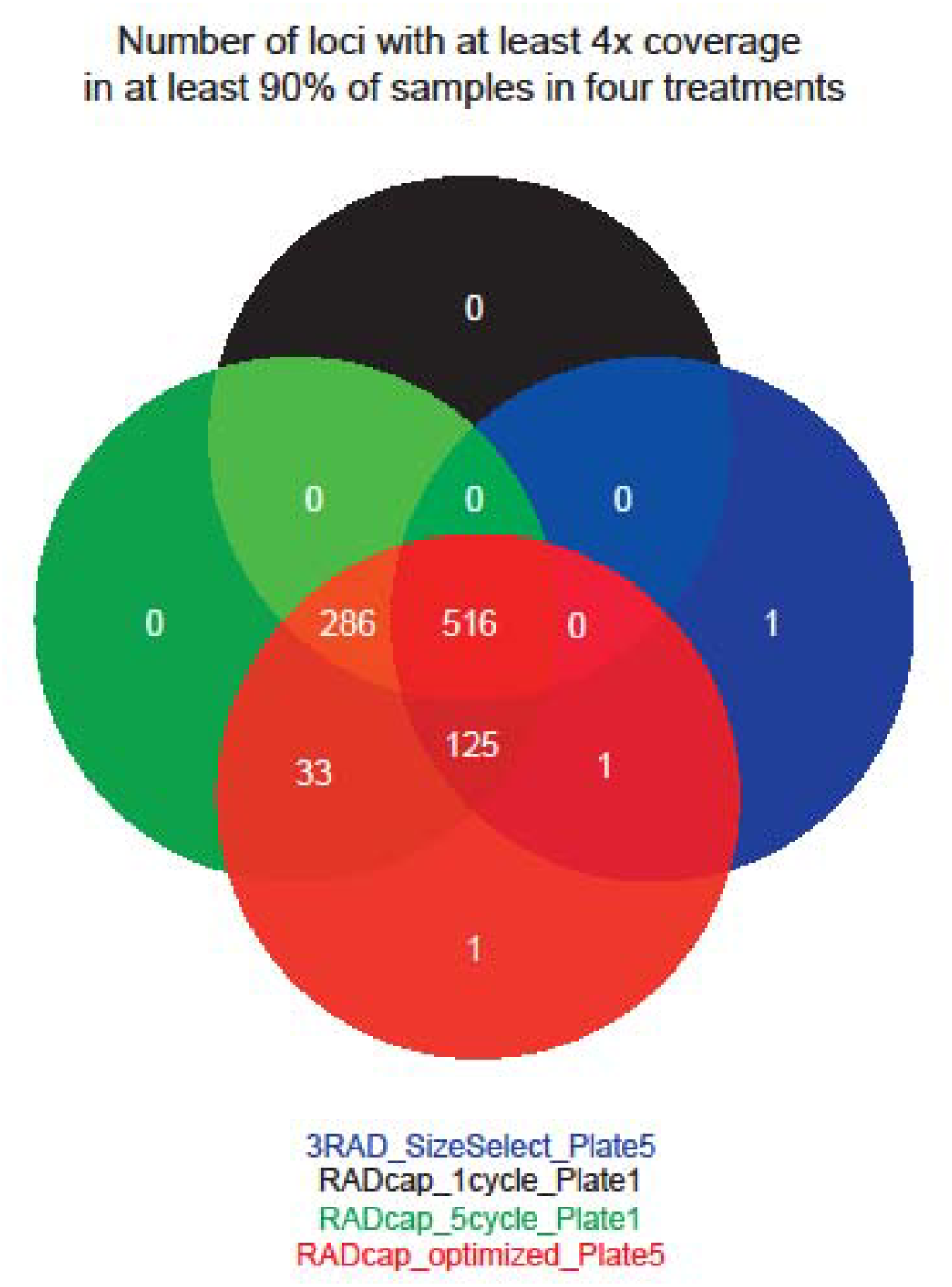
Venn diagram of the number of loci with at least 4x coverage shared across at least 90% (86) of samples in a single plate of the one-primer, two-primer, optimized, and size-selected treatments. Almost all of the 964 loci were recovered in the optimized and 5-cycle treatments, while the fewest loci were recovered in the size-selected treatment. Over half of the loci were shared among all four treatments, and only 3.7% of loci were found in one or two treatments.

In both RADcap_optimized_Plate5 and RADcap_5cycle_Plate1, we recovered most of the 964 loci in most samples regardless of the sequencing depth (Figure 6). By comparison, in the 3RAD with size selection treatment, even samples with the largest number of reads did not include as many loci as these RADcap treatments. When we modeled a reduced number of reads over all samples for each locus in the RADcap_optimized_Plate5 treatment, we found that 20,000 to 30,000 reads were sufficient to capture all loci with at least 4x coverage, and 60,000 reads per sample were sufficient to achieve 10x coverage at all loci (Figure 7). To achieve 20x or higher coverage at all loci, we estimated that ≥100,000 reads per sample were required, although it may not be economical to sequence all loci at 20x coverage or higher (see Discussion).

**Figure 6:**
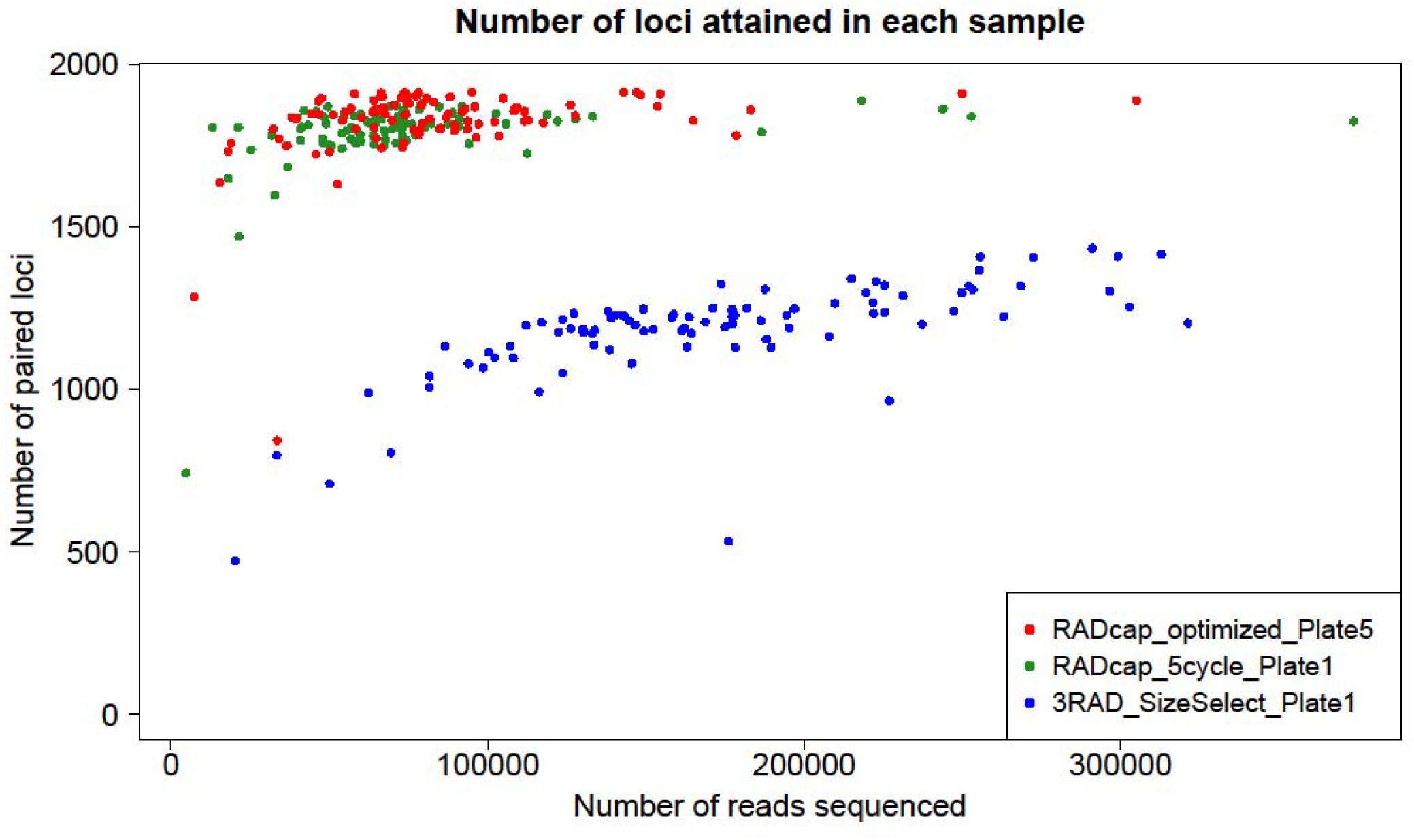
A scatterplot of the number of paired loci sequenced to ≥4x coverage in all samples and the number of reads sequenced for each sample. All samples in plates RADcap_optimized_Plate5, RADcap_5cycle_Plate1, and 3RAD_SizeSelect_Plate1 had between approximately 20,000 and 350,000 reads. Most samples in the optimized and two-primer treatments with at least 50,000 reads had all paired loci, whereas most size-selected samples, even those with 300,000 reads, did not have all paired loci.

**Figure 7:**
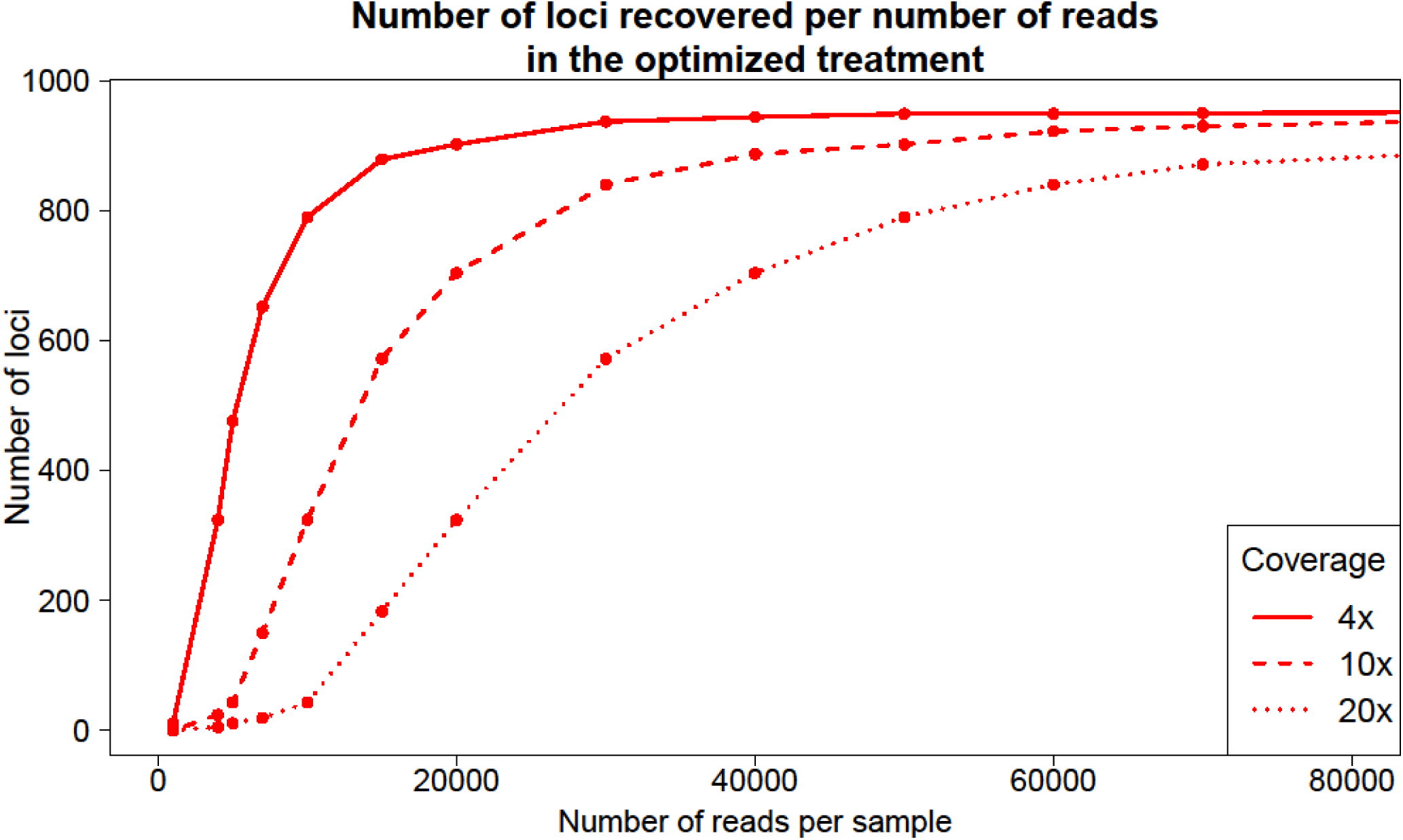
The number of loci that should be recovered at various read depths for a minimum coverage of 4x, 10x, and 20x. For 4x coverage, 30,000 reads is enough to sequence all loci, whereas, 60,000 reads per sample is required for 10x coverage. For 20x coverage, over 80,000 reads are required, and it may not be practical to sequence all loci at 20x coverage or higher.

### Error rate and Genetic Diversity of *Wisteria*

The amount of missing data in samples replicated within a treatment was effectively equal between 1-cycle and 5-cycle treatments (7.20% and 7.61%, respectively).

Because the optimized and size-selected treatments had the same samples and were filtered to have the same occupancy, we show the effect of estimating diversity with a smaller dataset. In the 80% occupancy matrices, we recovered 3744 SNPs in the optimized treatment and 2554 SNPs in the size-selected treatment. On average, two more samples were genotyped in the optimized treatment than size-selected treatment for each SNP (Table 3). The number of alleles number of effective alleles per SNP, Shannon’s information index, and F_IS_ were higher for size-selected samples than optimized samples. Although observed heterozygosity was the same, expected heterozygosity was higher for size-selected samples.

**Table 3:** The number of samples genotyped (N), number of alleles (Na), effective number of alleles (Ne), Shannon’s Information Index (I), observed (Ho) and expected (He) heterozygosity, and fixation index (F_IS_) as calculated in GenAlEx for plate 5 prepared via RADcap and 3RAD.

| Plate | N | Na | Ne | I | Ho | He | F<sub>IS</sub> |
| --- | --- | --- | --- | --- | --- | --- | --- |
| RADcap_optimized_Plate5 | 93.807 ±<br>0.061 | 2.014 ±<br>0.003 | 1.143 ±<br>0.004 | 0.169 ±<br>0.003 | 0.103 ±<br>0.003 | 0.095 ±<br>0.002 | 0.180 ±<br>0.007 |
| 3RAD_SizeSelect_Plate5 | 91.575 ±<br>0.106 | 2.018 ±<br>0.003 | 1.159 ±<br>0.005 | 0.184 ±<br>0.004 | 0.104 ±<br>0.004 | 0.104 ±<br>0.003 | 0.227 ±<br>0.008 |

## Discussion

Our overall goal was to develop a simple system that would efficiently sample a consistent portion of the genome from large numbers of individuals at low cost. Our optimized protocol achieves ≥4x coverage for ≥90% of samples at 99.8% of targeted loci with <187,000 reads per sample (Figure 4). This high sequencing depth means that the optimized protocol could achieve ≥4x coverage for >90% of 96 samples at >900 loci with only an average of 20,000 reads per sample (Figure 7). Furthermore, a single researcher was able to process >1000 DNA samples within a week and to sequence that pool with less than one-third of a NextSeq high output run (i.e., ~$1/sample in sequencing costs). RADcap performs exceedingly well and achieves our overall goal. It is well known, however, that 4x coverage will often lead to inaccurate genotypes and that deeper sequencing is needed for consistent and accurate genotyping (DePristo *et al*. 2011; Sims *et al*. 2014). Fortunately, RADcap is also sufficiently efficient that 10-20x coverage can be obtained for 90% complete matrices with affordable amounts of sequencing (Figure 7).

We discuss each major facet of our RADcap approach, the results we have obtained, and we note additional ways to implement and expand upon the RADcap technique (Table 4). Other groups with different goals have also combined RADseq with sequence capture (Jones & Good 2015), such as Suchan et al.’s (2015) use of RADseq fragments as baits. While completing this manuscript, a separate group published a similar method of sequence capture (Rapture; Ali *et al*. 2015), which shares the same general goal as our approach, and we discuss some similarities and differences between RADcap and Rapture.

**Table 4:** Major processes and reagents of RADcap, current costs, and potential improvements to reduce costs and/or increase throughput. Current costs of processes are calculated on a persample basis for the methods used herein and assume full 96-well plates. Current costs of major reagents are the batch cost (per project).

| Process | Reagent cost (\$US) | Alternatives and Potential Improvements |
| --- | --- | --- |
| Isolating DNA | 2 | Speed-bead based DNA isolation |
| Normalizing DNA | 0.2 | Robots, Sequal-Prep or similar |
| Digesting DNA & Ligating Adapters | 1 | Reduce amount of enzymes used |
| Single cycle degenerate iTru5 | 0.2 |  |
| Size Selection (SNP/locus discovery) | 0.1<sup>¶</sup> | Speed-beads or gel-cut |
| NextSeq PE150 | 2<sup>¶</sup> | HiSeq PE100 – PE150 |
| Sequence Capture | 1.2 | - |
| NextSeq PE75 | 1.2 |  |
| RADcap genotyping total per sample | 5.8 |  |
| **Major Reagents** |  |  |
| MYcroarray MYbaits | 2400* | Single-baits per locus, Increase number of projects pooled for baits |
| 3RAD Adapters | 370<sup>§</sup> | Purchase aliquots or share |
| iTru Primers | 345<sup>§</sup> | Purchase aliquots or share |

<sup>¶</sup>Excluded in the total genotyping cost per sample; \*included or <sup>§</sup>excluded in per sample cost

### Initial 3RAD SNP Discovery and Bait Design

RADcap is not limited to 1000 loci. Our initial goal was to obtain SNP discovery data for >2000 polymorphic loci, to design baits for the best ~1000 loci, and to retain a set of ~500 loci after data filtering that consistently produced high-quality genotypes. Stacks performed well for the task of identifying SNPs and polymorphic loci – using four individuals for initial SNP discovery yielded the desired number of polymorphic loci (2573). However, we recognize that using only four individuals is *not* ideal because it limits the ability to identify polymorphic loci that result from biological variation instead of errors. In addition, polymorphic loci with rare alleles were likely not attained in this small sample due to ascertainment bias (Clark *et al*. 2005; Nielsen 2000). For future RADcap projects, we will use 16-96 individuals for SNP discovery. One constraint of our current approach is that the pilot-scale 3RAD experiment requires a significant amount of time to complete, including a potential queue for the Illumina machine and several weeks to synthesize baits. However, if a genome sequence is available for the focal organisms, the genome could be digested *in silico* and loci with mapped SNPs could be used for bait design.

### Random Tagging at the i5 Index Position Allows Removal of PCR Duplicates

We were motivated to develop a system to remove PCR duplicates from RADseq libraries to achieve several goals: 1) reduce artificial confidence in genotypes resulting from undetected duplicates in ddRAD-type data, 2) satisfy the assumptions for data input to probabilistic base callers such as GATK, and 3) develop a general approach to identify PCR duplicates that would be easy to implement and optimize in a variety of experimental conditions. To achieve these goals, we implemented a new iTru5 primer with random i5 index sequences (iTru5-8N) for the Adapterama system (Glenn *et al*. 2016a; Glenn *et al*. 2016b; Glenn *et al*. 2016c) and a single-cycle of strand extension to incorporate iTru5 sequences into new library strands. The advantages of our approach include its: a) low-cost, b) simplicity, and c) freedom from requiring changes to standard Illumina sequencing or data processing protocols.

We used the iTru5-8N tag to successfully remove PCR duplicates from our data using new additions to the Stacks codebase, which is desirable for methods such as Stacks that rely on coverage to determine whether a variant is real or an artifact of PCR or sequencing (Casbon *et al*. 2011). Although there are 4^8^ = 65,536 possible iTru5-8N sequence tags for each locus, false duplicates (i.e., independent DNA molecules with the same iTru5-8N sequence tag) will be encountered at much lower coverage (Schweyen *et al*. 2014), similar to how a relatively small group of people is likely to have a pair that share the same birthday (McKinney 1966). The number of iTru5-8N sequence tags that we used to identify duplicates is much larger than tag pools used in the past (e.g., Schweyen *et al*. 2014; Tin *et al*. 2015), allowing more than sufficient depth of coverage after duplicate removal. The approach we created does not require researchers to anneal complementary oligos within a large pool of oligos containing degenerate tags (cf. Schweyen *et al*. 2014), which will produce a preponderance of double-stranded adapters with mismatches.

While our approach is simple and powerful, it comes with several limitations. First, there is an upper limit of 65,536 possible iTru5-8N sequence tags, thus the method we implemented in Stacks uses the iTru5-8N sequence, plus the sequences to which any given iTru5-8N sequence tag is incorporated to define duplicates, otherwise only 65,536 reads would be retained from any library. Second, it is critical to use conditions that promote high efficiency of first strand synthesis (i.e., our optimized treatment, and not the one-primer, two-primer, or low-template treatments) to avoid high levels of PCR duplication. Finally, Stacks is currently the only software that has been created and optimized to remove duplicates from these types of data.

Casbon *et al*. (2011) found that reduced amounts of template molecules going into PCR increased the rate of PCR duplication. In contrast, our treatment with the least amount of input DNA, RADcap_Low_Template_Plate1, had fewer duplicates than RADcap_1cycle_Plate1.

Thus, the specific conditions used for first strand extension are critical to producing a diverse RAD library and can be even more important than the amount of template used. Comparing the number of duplicates in the one-primer and two-primer treatments further illustrates the importance of the conditions used for first strand extension. The strand extension conditions as well as the starting DNA quality and quantity in plates 1-4 of the one-primer and two-primer treatments were identical, thus the reduced number of duplicates identified in the two-primer treatment relative to the one-primer treatment is due to hidden duplicates in the two-primer treatment. The low-level of duplicates in the size-selected and optimized protocols (which have only one cycle of first strand extension) demonstrates that high levels of duplicates are not inevitable, and that careful optimization of reaction conditions can keep duplicates to quite low proportions. However, the only way to know what percentage of the reads are duplicates is to implement a strategy to detect duplicates, which also facilitates their removal. Thus, tagging and removing duplicates is prudent for all RADcap and RADseq experiments.

### Optimizing RADcap Efficiency

Our modifications of the 1-cycle treatment to the optimized (also single-primer) treatment included increasing the PCR volume and adding locus-specific bait blockers for the few loci that were over-abundant in the first set of RADcap reads. These modifications decreased the variation in coverage among loci and increased the number of loci we recovered.

Although the bait-blockers produced an effect (the percentage of reads attributed to the blocked loci decreased from 14.7% in the 1-cycle treatment and 8.1% in the 5-cycle treatment to 3.6% in the optimized treatment), the size of this effect was modest. Thus, we surmise that increasing the PCR volume used for first-strand synthesis was far more important. Unfortunately, because we tested the one-primer and two-primer treatments on plates 1-4 while we tested the increased PCR volume using plate 5, it is unclear whether the initial library quality of plate 5 was higher than plates 1-4 or if the increased volume of the PCR reaction decreased the rate of PCR duplication. We do not expect or have evidence that the DNA varied in any significant way among plates, but we cannot rule out that possibility. Thus, although the optimized treatment conditions appear to have produced a very significant reduction in duplicates, additional experiments are necessary to definitively reach that conclusion.

### RADcap Captures Nearly All Loci Targeted

We sequenced most of the targeted loci in RADcap_optimized_Plate5 and only slightly fewer loci in RADcap_5cycle_Plate1 (Figure 4). The 99.8% overlap between RADcap_optimized_Plate5 and RADcap_5cycle_Plate1 illustrates the strength of using sequence capture to collect RAD loci: we were able to recover most of the same loci across at least 90% of 192 samples that we prepared several weeks apart.

Although RADseq protocols reduce the genome being studied, the number of loci in a typical analysis can still be in the tens of thousands. Our RADcap method allows for sequencing efforts to be further focused, permitting higher levels of multiplexing and locus recovery in an experiment while avoiding some of the problems such as biases in PCR amplification, variation at restriction enzyme cut sites, and purification and size selection methods (DaCosta & Sorenson 2014; Gautier *et al*. 2013) that can occur with ddRAD or traditional RADseq experiments implemented at the same scale.

### RADcap Produces Dense Matrices

We found that the probabilistic base-caller, GATK, worked well on these data, recovering and retaining large numbers of loci and SNPs at 50%, 80%, and 90% matrix occupancy (Figure 5). The number of loci and SNPs called by GATK follows predictable patterns within and among datasets from all treatments. We selected GATK because it is in common use across a variety of genotyping studies and because it performs well for moderate- to large-scale data sets. However, GATK is unsuited to most ddRAD data sets because of read duplication, thus our system for removing duplicates was critical for meeting the assumptions of GATK. Obviously, we could have used Stacks or other SNP-calling software packages (e.g. FreeBayes, pyRAD, SAMTools), and subsequent work will provide a detailed comparison among SNP-calling software packages.

### RADcap Adds Relatively Few Errors to Illumina Sequences

Errors in RADseq data derive from library preparation and sequencing methods. Errors introduced by PCR may be common in RADseq data because even extremely high fidelity DNA polymerases introduce errors on the order of 2.8×10^-7^ per nucleotide incorporated (KAPA, Boston, MA, USA). If these errors occurred in a single cycle of PCR, it would result in 4,683 errors in the 223 million reads in the present dataset. Because there can be many PCR cycles in RADseq library preparation, an incorrect base incorporated during early cycles be amplified to high coverage in the dataset. A much larger problem is the 0.1% substitution error rate made by Illumina machines (Glenn 2011), which results in an additional 28,544,000 incorrect bases in a dataset of 223 million PE64 reads. Even very small biases away from randomness in the Illumina sequencing error distribution can create significant downstream problems in data analysis. Decloning facilitated by the random iTru5-8N primer tags does not prevent PCR or sequencing errors, but the use of probabilistic base calling algorithms as implemented in GATK or other SNP-calling software can help reduce the likelihood of a base introduced by these errors from being called as a true variant.

### RADcap Works with Mixtures of Baits from Different Organisms Diluted to 1x Concentration

We used a bait set containing baits from two organisms – 2328 baits from our focal *Wisteria* groups and 2624 baits from an unrelated organism (kudzu; *Pueraria montana* var. *lobata*). As a result, baits for both species were present in all captures, despite DNA from only one species being present in any given capture. We also synthesized fewer than the maximum allowed number of baits (i.e. 4,952 baits in synthesis scale meant for up to 20,000 baits). The large number of loci we captured and the dense genotype matrices we created suggest that there was no meaningful interference from the additional baits during sequence capture and that the concentration of baits we applied to each sample pool was sufficient.

By mixing baits for two different projects, we were able to reduce baits costs by 50% for each project. The MYbaits-1 synthesis allows ~20,000 baits and the smallest synthesis scale is sufficient for 12 captures. If fewer than 20,000 baits are needed, then the concentration of baits is increased proportionately (e.g., 20,000 baits at 1x or 10,000 baits at 2x). Baits can also be divided among multiple projects. For example, researchers could pool four different projects, each with 1000 baits, into a single MYbaits-1 synthesis to achieve baits at 5x normal concentration. This would mean workers could achieve 60 (12×5) captures of 96 samples, rather than 12. Divided evenly, this would allow 15×96 (1440) samples from each of the 4 projects to be enriched when purchasing the smallest possible single MYbaits-1 synthesis. Obviously, different numbers of taxa with varying numbers of samples could be accommodated (e.g., 5,000 samples from one taxon and 760 samples from the other three; see Heyduk *et al*. (2016) for additional examples). This flexibility creates many opportunities for collecting RADcap data at low cost.

### Comparison of RADcap to Rapture

Rapture is a similar, enrichment-based, RAD sequencing approach that uses a two-step protocol to capture RADseq loci. In the first step, researchers digest, randomly shear, and ligate biotinylated adapters to DNA, which can then be separated from other genomic fragments with magnetic beads before library preparation continues. In the second step, similar to RADcap, custom library-specific baits are used to capture loci of interest. RADcap and Rapture are similar in that they require DNA isolation, restriction enzyme digests, ligation of adapters, pooling, clean-up, capture, and sequencing. Both methods are significant advances that increase the density and consistency of genotype matrices while simultaneously reducing costs for large-scale projects.

There are, however, significant differences in cost and flexibility between RADcap and Rapture as a result of RADcap’s integration with 3RAD and the Adapterama system. The 3RAD adapters require 8 phosphorylated oligos and 32 plain oligos to achieve 96 combinations (Glenn *et al*. 2016b), whereas Rapture requires 96 biotinylated oligos plus 96 phosphorylated oligos, making Rapture adapters 10 times more expensive ($370 for RADcap vs. $3750 for Rapture). Adding or switching to different enzymes in Rapture requires additional sets of adapters at $3750 per enzyme, whereas 3RAD facilitates the use of many different possible enzymes and combinations of enzymes (Glenn *et al*. 2016b) with fewer sets of interchangeable adapters at $370 per set. In addition, RADcap does not require any commercial library preparation kits, whereas Rapture makes use of commercial library preparation kits.

In addition to cost differences, duplicate detection has fewer false positives in RADcap than Rapture. Rapture detects duplicates based on the starting position of Read 2, which may be anywhere along ~500 bases (following shearing and size selection). Thus, RADcap has 65,536 tags, whereas Rapture has ~500. Additionally, dual digest RADcap increases coverage at both ends of the library molecule, and we show that fewer reads per sample are required to achieve the same coverage with RADcap than Rapture (20,000 versus 50,000 for at least 4x coverage, respectively). On the other hand, Rapture’s use of random shearing increases the length of the genomic region that is sequenced, which may be an advantage worth the trade-off in decreased coverage, depending upon the goals of the project.

### Future Improvements and Extensions

RADcap (and Rapture) open the door to a variety of additional research opportunities. One of the most important is the option of using the capture baits from RAD loci on randomly-sheared genomic libraries (i.e., standard genomic libraries). Such work will facilitate direct comparisons between RAD loci and other loci commonly used for sequence capture (exons, UCEs, anchored loci, etc.). Although preparing randomly sheared genomic libraries for RADcap increases the cost per sample, it will allow: 1) assembling contigs at captured loci so that more sequence is available to facilitate a) ortholog identification in other species, b) identifying additional linked SNPs, c) phasing SNPs within captured loci, and d) better understanding of the sequence context for the RADcap loci; 2) investigating rates of divergence at restriction sites; 3) collecting RAD loci from samples with deeper divergences than is feasible with restriction sites (i.e., for phylogenetics), and 4) using PHYLUCE (Faircloth 2015) and other analytical tools that have been developed for UCEs and other sequence capture systems. Capture baits also facilitate using RADseq for degraded and contaminated samples (cf. Graham *et al*. 2015) and focusing on microsatellite loci present in RADseq libraries, either through the use of locus-specific baits that target the flanking regions or via generic baits to the repeats (cf. Glenn & Schable 2005). Thus, the baits identified with RADcap will serve a variety of purposes in future work.

Given our high efficiency with two baits per locus, future work should investigate the efficiency of capturing loci with a single bait per locus where we suspect the success rate will remain high because of the reduced diversity in RAD libraries. We have presented datasets and results obtained from the first sets of samples we have conducted with RADcap. Although we are bullish on the future of RADcap, we expect researchers will explore and discover how changes to the protocols (often unintentional), as well as their use on different organisms, impact the outcome.

## Summary

We present a novel protocol to cheaply sequence a specific set of hundreds to thousands of loci in hundreds to thousands of samples. We demonstrate a generalizable method for identifying PCR duplicates in Illumina libraries, even double-digest, RADseq-type libraries. We also show that it is possible to reduce PCR duplicates to 5% of the total library, to routinely achieve >80% on-target reads, and to achieve dense matrices of genotypes from hundreds of individuals. Our method is sufficiently efficient that researchers can choose to sequence less deeply to achieve commonly observed coverage in RADseq-type datasets, or they can affordably sequence much more deeply to obtain high-quality genotypes. Molecular ecology research is filled with choices about where to spend limited resources (time and money). We strongly recommend that researchers adopt methods that yield high coverage and dense matrices of high-confidence genotypes, and we hope that RADcap allows other scientists to obtain high-quality data and make more robust conclusions about their study systems.

## Acknowledgements

We thank Todd Pierson, Kerin Bentley, and Natalia Bayona. This work was supported by grants DEB-1242260 and DEB-1146440 from the U.S. National Science Foundation and the U.S. National Science Foundation Partnership for International Research and Education (PIRE) program (OISE 0730218). This study was also supported in part by resources and technical expertise from the Georgia Advanced Computing Resource Center, a partnership between the University of Georgia’s Office of the Vice President for Research and Office of the Vice President for Information Technology.

## Author Contributions

Conceived RADcap: TCG, BCF; conceived *Wisteria* project: SLH, RM; conceived and implemented decloning software: JMC; prepared samples: SLH, TJK; designed baits: AD; analyzed data: SLH, BCF; wrote paper: SLH, BCF, TCG, RM; contributed funding and other resources: TCG, RM; all authors edited and approved of the final version.

## Data Accessibility

Raw Reads will be at NCBI SRA: xxxxxxxx.

## Literature Cited

Ali OA, O'Rourke SM, Amish SJ, et al. (2015) RAD Capture (Rapture): Flexible and efficient sequence-based genotyping. bioRxiv, 028837.

Altshuler D, Pollara VJ, Cowles CR, et al. (2000) An SNP map of the human genome generated by reduced representation shotgun sequencing. Nature 407, 513-516.

Andrews KR, Good JM, Miller MR, Luikart G, Hohenlohe PA (2016) Harnessing the power of RADseq for ecological and evolutionary genomics. Nat Rev Genet 17, 81-92.

Andrews KR, Hohenlohe PA, Miller MR, et al. (2014) Trade-offs and utility of alternative RADseq methods: Reply to Puritz et al. 2014. Molecular ecology 23, 5943-5946.

Baird NA, Etter PD, Atwood TS, et al. (2008) Rapid SNP Discovery and Genetic Mapping Using Sequenced RAD Markers. PLoS ONE 3, e3376.

Bansal V, Harismendy O, Tewhey R, et al. (2010) Accurate detection and genotyping of SNPs utilizing population sequencing data. Genome research 20, 537-545.

Bi K, Vanderpool D, Singhal S, et al. (2012) Transcriptome-based exon capture enables highly cost-effective comparative genomic data collection at moderate evolutionary scales. BMC Genomics 13, 1-14.

Boratyn GM, Camacho C, Cooper PS, et al. (2013) BLAST: a more efficient report with usability improvements. Nucleic Acids Research 41, W29-W33.

Cao H, Wu J, Wang Y, et al. (2013) An Integrated Tool to Study MHC Region: Accurate SNV Detection and HLA Genes Typing in Human MHC Region Using Targeted High-Throughput Sequencing. PLoS ONE 8, e69388.

Cariou M, Duret L, Charlat S (2013) Is RAD-seq suitable for phylogenetic inference? An in silico assessment and optimization. Ecology and Evolution 3, 846-852.

Casbon JA, Osborne RJ, Brenner S, Lichtenstein CP (2011) A method for counting PCR template molecules with application to next-generation sequencing. Nucleic Acids Research 39.

Catchen J, Hohenlohe PA, Bassham S, Amores A, Cresko WA (2013) Stacks: an analysis tool set for population genomics. Molecular ecology 22, 3124-3140.

Catchen JM, Amores A, Hohenlohe P, Cresko W, Postlethwait JH (2011) Stacks: Building and Genotyping Loci De Novo From Short-Read Sequences. G3-Genes Genomes Genetics 1, 171-182.

Chen H, Boutros PC (2011) VennDiagram: a package for the generation of highly-customizable Venn and Euler diagrams in R. BMC bioinformatics 12, 35.

Clark AG, Hubisz MJ, Bustamante CD, Williamson SH, Nielsen R (2005) Ascertainment bias in studies of human genome-wide polymorphism. Genome research 15, 1496-1502.

Craig DW, Pearson JV, Szelinger S, et al. (2008) Identification of genetic variants using bar-coded multiplexed sequencing. Nature Methods 5, 887-893.

DaCosta JM, Sorenson MD (2014) Amplification Biases and Consistent Recovery of Loci in a Double-Digest RAD-seq Protocol. PLoS ONE 9, e106713.

Danecek P, Auton A, Abecasis G, et al. (2011) The variant call format and VCFtools. Bioinformatics 27, 2156-2158.

Davey JL, Blaxter MW (2010) RADSeq: next-generation population genetics. Briefings in Functional Genomics 9, 416-423.

Davey JW, Hohenlohe PA, Etter PD, et al. (2011) Genome-wide genetic marker discovery and genotyping using next-generation sequencing. Nature Reviews Genetics 12, 499-510.

DePristo MA, Banks E, Poplin R, et al. (2011) A framework for variation discovery and genotyping using next-generation DNA sequencing data. Nat Genet 43, 491-498.

Faircloth BC (2015) PHYLUCE is a software package for the analysis of conserved genomic loci. Bioinformatics.

Faircloth BC, McCormack JE, Crawford NG, et al. (2012) Ultraconserved Elements Anchor Thousands of Genetic Markers Spanning Multiple Evolutionary Timescales. Systematic biology 61, 717-726.

Fountain ED, Pauli JN, Reid BN, Palsbøll PJ, Peery MZ (2016) Finding the right coverage: the impact of coverage and sequence quality on single nucleotide polymorphism genotyping error rates. Molecular Ecology Resources, n/a-n/a.

Gautier M, Gharbi K, Cezard T, et al. (2013) The effect of RAD allele dropout on the estimation of genetic variation within and between populations. Molecular ecology 22, 3165-3178.

Glenn TC (2011) Field guide to next-generation DNA sequencers. Molecular Ecology Resources 11, 759-769.

Glenn TC, Faircloth BC, Nilsen R, et al. (2016a) Adapterama I: Universal stubs and primers for thousands of dual-indexed Illumina libraries. in prep.

Glenn TC, Pierson T, Kieran TJ, et al. (2016b) Adapterama III: Quadruple-indexed triple-enzyme RADseq libraries from picograms of DNA (3RAD). in prep.

Glenn TC, Pierson TW, Kieran TJ, et al. (2016c) Adapterama II: Universal amplicon sequencing on Illumina platforms (TaggiMatrix). in prep.

Glenn TC, Schable NA (2005) Isolating Microsatellite DNA Loci. In: Methods in Enzymology, pp. 202-222. Academic Press.

Gnirke A, Melnikov A, Maguire J, et al. (2009) Solution hybrid selection with ultra-long oligonucleotides for massively parallel targeted sequencing. Nature Biotechnology 27, 182-189.

Gordon A, Hannon G (2010) Fastx-toolkit. FASTQ/A short-reads preprocessing tools (unpublished) http://hannonlab.cshl.edu/fastx_toolkit.

Graham CF, Glenn TC, McArthur AG, et al. (2015) Impacts of degraded DNA on restriction enzyme associated DNA sequencing (RADSeq). Molecular Ecology Resources.

Harvey MG, Smith BT, Glenn TC, Faircloth BC, Brumfield RT (2013) Sequence capture versus restriction site associated DNA sequencing for phylogeography. arXiv preprint arXiv:1312.6439.

Heyduk K, Trapnell DW, Barrett CF, Leebens-Mack J (2016) Phylogenomic analyses of species relationships in the genus Sabal (Arecaceae) using targeted sequence capture. Biological Journal of the Linnean Society 117, 106-120.

Hiatt JB, Pritchard CC, Salipante SJ, O'Roak BJ, Shendure J (2013) Single molecule molecular inversion probes for targeted, high-accuracy detection of low-frequency variation. Genome research 23, 843-854.

Jabara CB, Jones CD, Roach J, Anderson JA, Swanstrom R (2011) Accurate sampling and deep sequencing of the HIV-1 protease gene using a Primer ID. Proc Natl Acad Sci USA 108, 20166-20171.

Jones MR, Good JM (2015) Targeted capture in evolutionary and ecological genomics. Molecular ecology.

Kenny EM, Cormican P, Gilks WP, et al. (2010) Multiplex Target Enrichment Using DNA Indexing for Ultra-High Throughput SNP Detection. DNA Research.

Kivioja T, Vaharautio A, Karlsson K, et al. (2012) Counting absolute numbers of molecules using unique molecular identifiers. Nat Meth 9, 72-74.

Lemmon AR, Emme SA, Lemmon EM (2012) Anchored Hybrid Enrichment for Massively High-Throughput Phylogenomics. Systematic biology 61, 727-744.

Li C, Hofreiter M, Straube N, Corrigan S, Naylor GJ (2013) Capturing protein-coding genes across highly divergent species. Biotechniques 54, 321-326.

Li H, Durbin R (2009) Fast and accurate short read alignment with Burrows-Wheeler transform. Bioinformatics 25, 1754-1760.

Li H, Handsaker B, Wysoker A, et al. (2009) The Sequence Alignment/Map format and SAMtools. Bioinformatics 25, 2078-2079.

Lischer HEL, Excoffier L (2012) PGDSpider: an automated data conversion tool for connecting population genetics and genomics programs. Bioinformatics 28, 298-299.

Mastretta-Yanes A, Arrigo N, Alvarez N, et al. (2015) Restriction site-associated DNA sequencing, genotyping error estimation and de novo assembly optimization for population genetic inference. Molecular Ecology Resources 15, 28-41.

McKenna A, Hanna M, Banks E, et al. (2010) The Genome Analysis Toolkit: a MapReduce framework for analyzing next-generation DNA sequencing data. Genome Res 20, 1297-1303.

McKinney EH (1966) Generalized Birthday Problem. The American Mathematical Monthly 73, 385-387.

Miller MR, Dunham JP, Amores A, Cresko WA, Johnson EA (2007) Rapid and cost-effective polymorphism identification and genotyping using restriction site associated DNA (RAD) markers. Genome research 17, 240-248.

Miner BE, Stöger RJ, Burden AF, Laird CD, Hansen RS (2004) Molecular barcodes detect redundancy and contamination in hairpin-bisulfite PCR. Nucleic Acids Research 32, e135-e135.

Nielsen R (2000) Estimation of population parameters and recombination rates from single nucleotide polymorphisms. Genetics 154, 931-942.

Novaes E, Drost DR, Farmerie WG, et al. (2008) High-throughput gene and SNP discovery in Eucalyptus grandis, an uncharacterized genome. BMC Genomics 9, 1-14.

Okou DT, Steinberg KM, Middle C, et al. (2007) Microarray-based genomic selection for high-throughput resequencing. Nature Methods 4, 907-909.

Pante E, Abdelkrim J, Viricel A, et al. (2015) Use of RAD sequencing for delimiting species. Heredity 114, 450-459.

Peakall R, Smouse PE (2006) GENALEX 6: genetic analysis in Excel. Population genetic software for teaching and research. Molecular Ecology Notes 6, 288-295.

Peterson BK, Weber JN, Kay EH, Fisher HS, Hoekstra HE (2012) Double Digest RADseq: An Inexpensive Method for De Novo SNP Discovery and Genotyping in Model and Non-Model Species. PLoS ONE 7.

Pröll J, Danzer M, Stabentheiner S, et al. (2011) Sequence Capture and Next Generation Resequencing of the MHC Region Highlights Potential Transplantation Determinants in HLA Identical Haematopoietic Stem Cell Transplantation. DNA Research 18, 201-210.

Puritz JB, Matz MV, Toonen RJ, et al. (2014) Demystifying the RAD fad. Molecular ecology 23, 5937-5942.

Rabenstein H, Bangol B, Linke S, et al. (2015) Large fragment target enrichment and sequencing of the 4.4 Mb major histocompatibility complex by region-specific extraction and HiSeq 2500. Human Immunology 76, Supplement, 57.

Raposo do Amaral F, Neves LG, Resende MFR, Jr., et al. (2015) Ultraconserved Elements Sequencing as a Low-Cost Source of Complete Mitochondrial Genomes and Microsatellite Markers in Non-Model Amniotes. PLoS ONE 10, e0138446.

Rogers Y-H, Venter JC (2005) Genomics: Massively parallel sequencing. Nature 437, 326-327.

Rohland N, Reich D (2012) Cost-effective, high-throughput DNA sequencing libraries for multiplexed target capture. Genome research 22, 939-946.

Saintenac C, Jiang D, Akhunov ED (2011) Targeted analysis of nucleotide and copy number variation by exon capture in allotetraploid wheat genome. Genome Biology 12, R88-R88.

Schweyen H, Rozenberg A, Leese F (2014) Detection and Removal of PCR Duplicates in Population Genomic ddRAD Studies by Addition of a Degenerate Base Region (DBR) in Sequencing Adapters. Biological Bulletin 227, 146-160.

Shiroguchi K, Jia TZ, Sims PA, Xie XS (2012) Digital RNA sequencing minimizes sequence-dependent bias and amplification noise with optimized single-molecule barcodes. Proc Natl Acad Sci USA 109, 1347-1352.

Signorell A (2015) DescTools: Tools for descriptive statistics, p. R package version 0.99.15.

Sims D, Sudbery I, Ilott NE, Heger A, Ponting CP (2014) Sequencing depth and coverage: key considerations in genomic analyses. Nature Reviews Genetics 15, 121-132.

Smith E, Jepsen K, Khosroheidari M, et al. (2014) Biased estimates of clonal evolution and subclonal heterogeneity can arise from PCR duplicates in deep sequencing experiments. Genome Biology 15, 420.

Smouse PE, Peakall R (2012) GenAlEx 6.5: genetic analysis in Excel. Population genetic software for teaching and research—an update. Bioinformatics 28, 2537-2539.

Suchan T, Pitteloud C, Gerasimova N, et al. (2015) Hybridization capture using RAD probes (hyRAD), a new tool for performing genomic analyses on museum collection specimens. bioRxiv, 025551.

Tautz D, Ellegren H, Weigel D (2010) Next Generation Molecular Ecology. Molecular ecology 19, 1-3.

Tin MMY, Rheindt FE, Cros E, Mikheyev AS (2015) Degenerate adaptor sequences for detecting PCR duplicates in reduced representation sequencing data improve genotype calling accuracy. Molecular Ecology Resources 15, 329-336.

Trusty JL, Johnson KJ, Lockaby BG, Goertzen LR (2007) Bi-Parental Cytoplasmic DNA Inheritance in Wisteria (Fabaceae): Evidence from a Natural Experiment. Plant and Cell Physiology 48, 662-665.

Trusty JL, Lockaby BG, Zipperer WC, Goertzen LR (2008) Horticulture, hybrid cultivars and exotic plant invasion: a case study of Wisteria (Fabaceae). Botanical Journal of the Linnean Society 158, 593-601.

Valder P (1995) Wisterias: A comprehensive guide Timber Press, Potland, Oregon.

Wiedmann RT, Smith TP, Nonneman DJ (2008) SNP discovery in swine by reduced representation and high throughput pyrosequencing. BMC Genetics 9, 1-7.

Wilson EH (1916) The Wisterias of China and Japan. The Gardeners' Chronicle 1545, 61-62.

Wyman D (1949) The Wisterias. Arnoldia 9, 17-28.

